# Systematic characterization of gene function in a photosynthetic organism

**DOI:** 10.1101/2020.12.11.420950

**Authors:** Josep Vilarrasa-Blasi, Friedrich Fauser, Masayuki Onishi, Silvia Ramundo, Weronika Patena, Matthew Millican, Jacqueline Osaki, Charlotte Philp, Matthew Nemeth, Patrice A. Salomé, Xiaobo Li, Setsuko Wakao, Rick G. Kim, Yuval Kaye, Arthur R. Grossman, Krishna K. Niyogi, Sabeeha Merchant, Sean Cutler, Peter Walter, José R. Dinneny, Martin C. Jonikas, Robert E. Jinkerson

## Abstract

Photosynthetic organisms are essential for human life, yet most of their genes remain functionally uncharacterized. Single-celled photosynthetic model systems have the potential to accelerate our ability to connect genes to functions. Here, using a barcoded mutant library of the model eukaryotic alga *Chlamydomonas reinhardtii*, we determined the phenotypes of more than 58,000 mutants under more than 121 different environmental growth conditions and chemical treatments. 78% of genes are represented by at least one mutant that showed a phenotype, providing clues to the functions of thousands of genes. Mutant phenotypic profiles allow us to place known and previously uncharacterized genes into functional pathways such as DNA repair, photosynthesis, the CO_2_-concentrating mechanism, and ciliogenesis. We illustrate the value of this resource by validating novel phenotypes and gene functions, including the discovery of three novel components of a defense pathway that counteracts actin cytoskeleton inhibitors released by other organisms. The data also inform phenotype discovery in land plants: mutants in *Arabidopsis thaliana* genes exhibit similar phenotypes to those we observed in their Chlamydomonas homologs. We anticipate that this resource will guide the functional characterization of genes across the tree of life.

---

Major contributions to our understanding of gene functions in photosynthetic organisms have been made by studying microbial models, including the discovery and characterization of the Calvin-Benson-Bassham CO_2_ fixation cycle^1^ as well as the structures^2^, order^3^ and cloning^4^ of complexes in the photosynthetic electron transport chain. Advances in technology now provide opportunities for microbes to serve as powerful complements to land plants in the characterization of gene functions by enabling significantly higher experimental throughput^5^.

The single-celled green alga Chlamydomonas (*Chlamydomonas reinhardtii*) is a well-established model system for studies of key pathways including photosynthesis, primary metabolism, inter-organelle communication, and stress response^6^. Furthermore, amenability to microscopy and biochemical purifications have made Chlamydomonas a leading model system for studies of cilia^7–9^. Despite promising progress with the development of clustered regularly interspaced short palindromic repeats (CRISPR)-based reagents to generate targeted mutants^10,11^, low editing efficiencies currently prevent large scale CRISPR sgRNA library screens in Chlamydomonas. The recent generation of a barcoded Chlamydomonas mutant collection facilitates the study of individual genes and enables forward genetic screens^12^. In the present work, we leverage the amenability of Chlamydomonas for high-throughput methods to connect genotypes to phenotypes on a massive scale, allowing placement of genes into pathways and discovery of conserved gene functions in land plants.

## Systematic genome-scale phenotyping

To connect genotypes to phenotypes, we measured the growth of 58,101 Chlamydomonas mutants representing 14,695 genes (83% of all genes encoded in the Chlamydomonas genome, based on the current genome annotation, v5.6) under 121 environmental and chemical stress conditions (both control and experimental conditions are given in Table S1 and Table S2). We pooled the entire Chlamydomonas mutant collection and used molecular barcodes to quantify the relative abundance of each mutant after competitive growth (Fig. 1a-f). Growth conditions included heterotrophic, mixotrophic, and photoautotrophic growth under different photon flux densities and CO_2_ concentrations, as well as abiotic stress conditions such as various pH and temperatures. We also subjected the library to known chemical stressors, including DNA damaging agents, reactive oxygen species, antimicrobial drugs such as paromomycin and spectinomycin, as well as the actin-depolymerizing drug latrunculin B (LatB). To further expand our knowledge of chemical stressors in photosynthetic organisms, we identified 1,222 small molecules from the Library of AcTive Compounds on Arabidopsis (LATCA)^13^ that negatively influence Chlamydomonas growth (Extended Data Fig. 1, Table S3, Extended Data File 1), and performed competitive growth experiments in the presence of 52 of the most potent compounds. Taken together, this effort represents, to the best of our knowledge, the largest genotype-by-phenotype dataset to date for any photosynthetic organism, with 62 million data points (Table S4).

**Figure 1.**
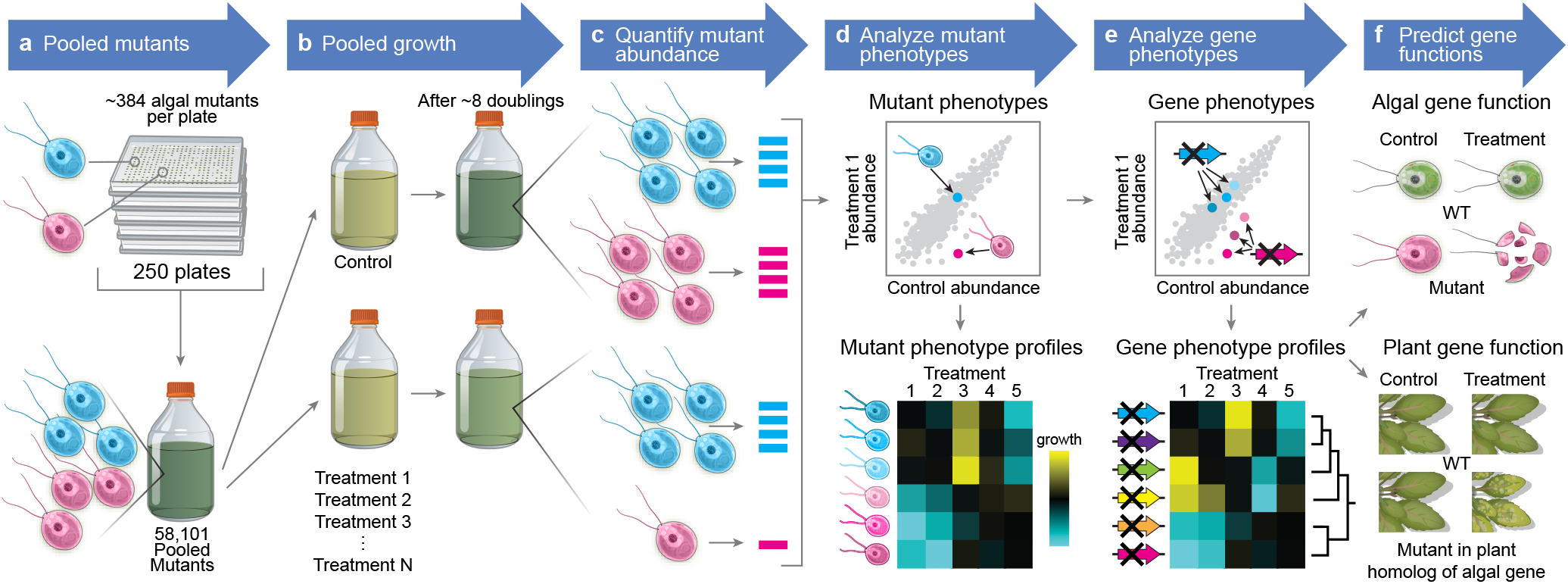
We developed a platform for genotype-phenotype discovery in a unicellular photosynthetic eukaryote. **a**. The Chlamydomonas mutant library was pooled and used to prepare a homogenous liquid starting culture of 58,101 mutants. **b**. Aliquots of the starting culture were used to inoculate pooled growth experiments to assess the fitness of each mutant under a variety of environmental and chemical stress treatments. **c**. The relative abundance of each mutant was quantified via PCR-based amplification of individual mutant barcodes and subsequent Illumina sequencing. **d**. Mutants negatively affected by the treatment have a lower barcode read count compared to the control. **e**. Many genes were represented by multiple mutants, which allowed the identification of high-confidence gene phenotypes. We then clustered genes based on their phenotypic profile to place genes into pathways and predict the functions of previously uncharacterized genes. **f**. The data predict gene function in Chlamydomonas and land plants.

## Mutants show genotype-phenotype specificity and enrichment of expected functions

To identify mutants with growth defects or enhancements due to a specific treatment, we compared the abundance of each mutant after growth under the treatment condition to its abundance after growth under a control condition (Fig. 2a). We called this comparison a screen, and the ratio of these abundances the mutant phenotype (Fig. 2b,c). Mutant phenotypes were reproducible between independent replicates of a screen (Fig. 2c,d).

**Figure 2.**
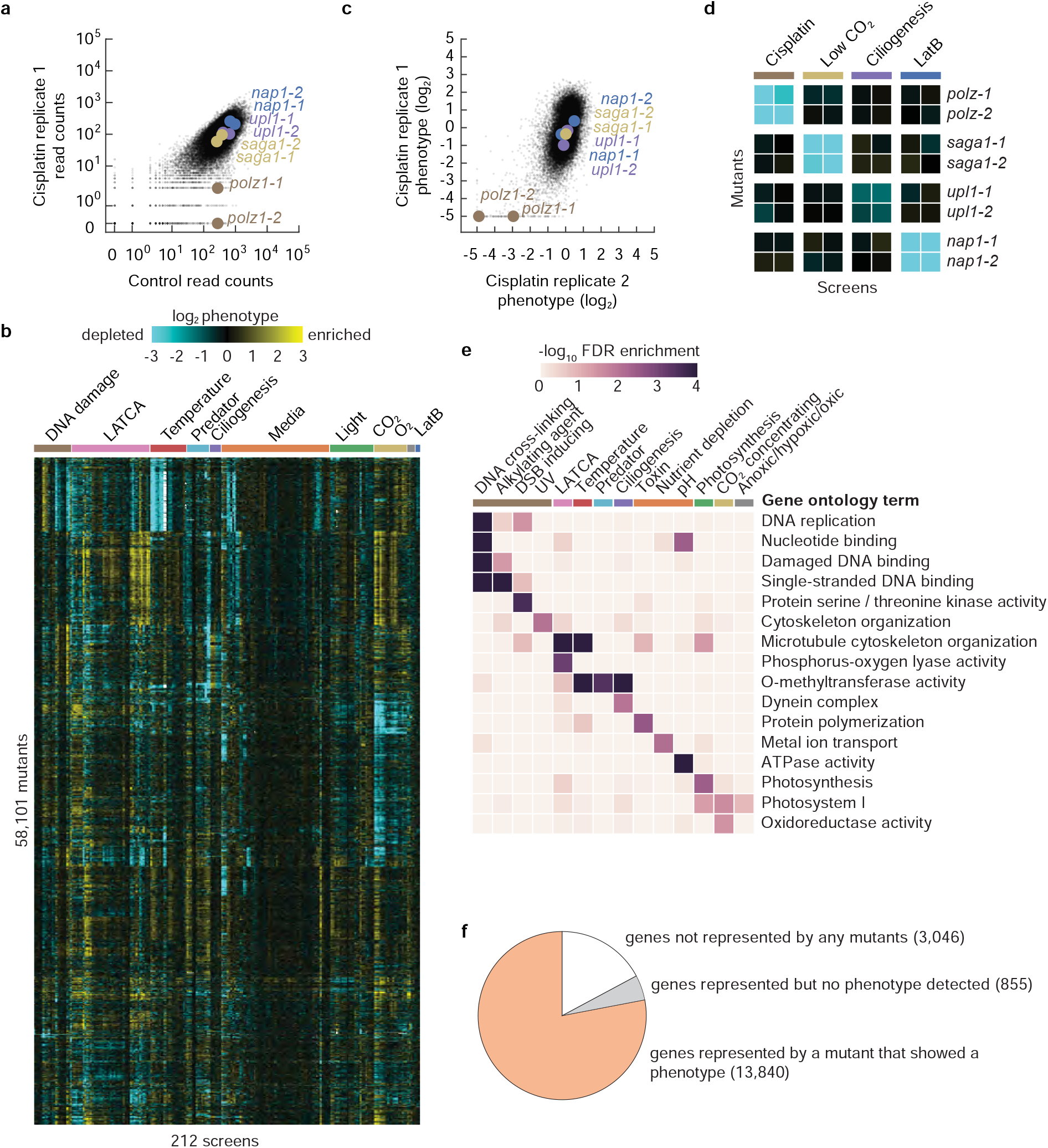
We determined the fitness of 58,101 Chlamydomonas mutants under 121 growth conditions. **a**. The phenotype of each mutant was determined by comparing its molecular barcode read count under a treatment and control condition. As an example, results from a screen using the drug cisplatin are shown. **b**. The typical reproducibility is illustrated with two replicate cisplatin screens. **c**. A hierarchically clustered heatmap shows the phenotype [log2(treatment reads/control reads)] of mutants across 212 screens representing 121 growth conditions. **d**. Mutants show screen-specific phenotypes. **e**. Gene Ontology term analysis reveals enrichment of biological functions associated with specific screens. **f**. Most genes are represented by at least one mutant that shows a phenotype in at least one treatment condition.

Individual mutants exhibited genotype-phenotype specificity. For example, mutants disrupted in the DNA repair gene *POLYMERASE ZETA* (*POLZ*, encoded by Cre09.g387400) exhibited growth defects in the presence of the DNA crosslinker cisplatin, and these mutants did not show growth defects in unrelated screens (Fig. 2d). We observed similar genotype-phenotype specificity for other genes and phenotypes including sensitivity to low CO_2_, ciliogenesis, and latrunculin B (LatB) sensitivity (Fig. 2d).

In many screens, mutants that exhibited phenotypes were enriched for disruptions in genes with expected function. 46 out of 223 screens, show at least one enriched (FDR <0.05) Gene Ontology (GO)^14^ term associated with mutants (Fig. 2e, Extended Data Fig. 2, Table S5). These enriched GO terms corresponded to functions known to be required for survival under the respective treatments. For example, screens with DNA-damaging agents resulted in GO term enrichments such as “DNA replication,” “Nucleotide binding,” or “Damaged DNA binding.” These GO term enrichments suggest that the phenotypic screens are correctly identifying genes required for growth under the corresponding stresses.

13,840 genes (78% of all Chlamydomonas genes) are represented by one or more mutant alleles that showed a phenotype (decreased abundance below our detection limit) in at least one screen. While a lone mutant showing a phenotype is not sufficient evidence to conclusively establish a gene-phenotype relationship, we anticipate that these data will be useful to the research community in at least three ways: first, they can help prioritize the characterization of candidate genes identified by other means, such as transcriptomics or protein-protein interactions. Second, they facilitate the generation of hypotheses about the functions of poorly-characterized genes. Third, they enable prioritization of available mutant alleles for further study. The genotype-phenotype specificity of individual mutants and the enrichment of expected functions suggest that our data can serve as a guide for understanding the functions of thousands of poorly-characterized genes.

## High-confidence, novel gene-phenotype relationships

The availability of multiple independent mutant alleles for individual genes allowed us to identify high-confidence gene-phenotype relationships. When multiple independent mutant alleles for the same gene show the same phenotype, the confidence in a gene lesion-phenotype relationship increases because it is less likely that the phenotype is due to a mutation elsewhere in the genome, or that there was an error in mapping of the mutation^12^. Using a statistical framework that leverages multiple independent mutations in the same gene (see Methods), we identified 1,636 high-confidence (FDR <0.3) gene-phenotype relationships involving 684 genes (Fig. 3a, Table S6, Table S7), including hundreds of genes with no functional annotation.

**Figure 3.**
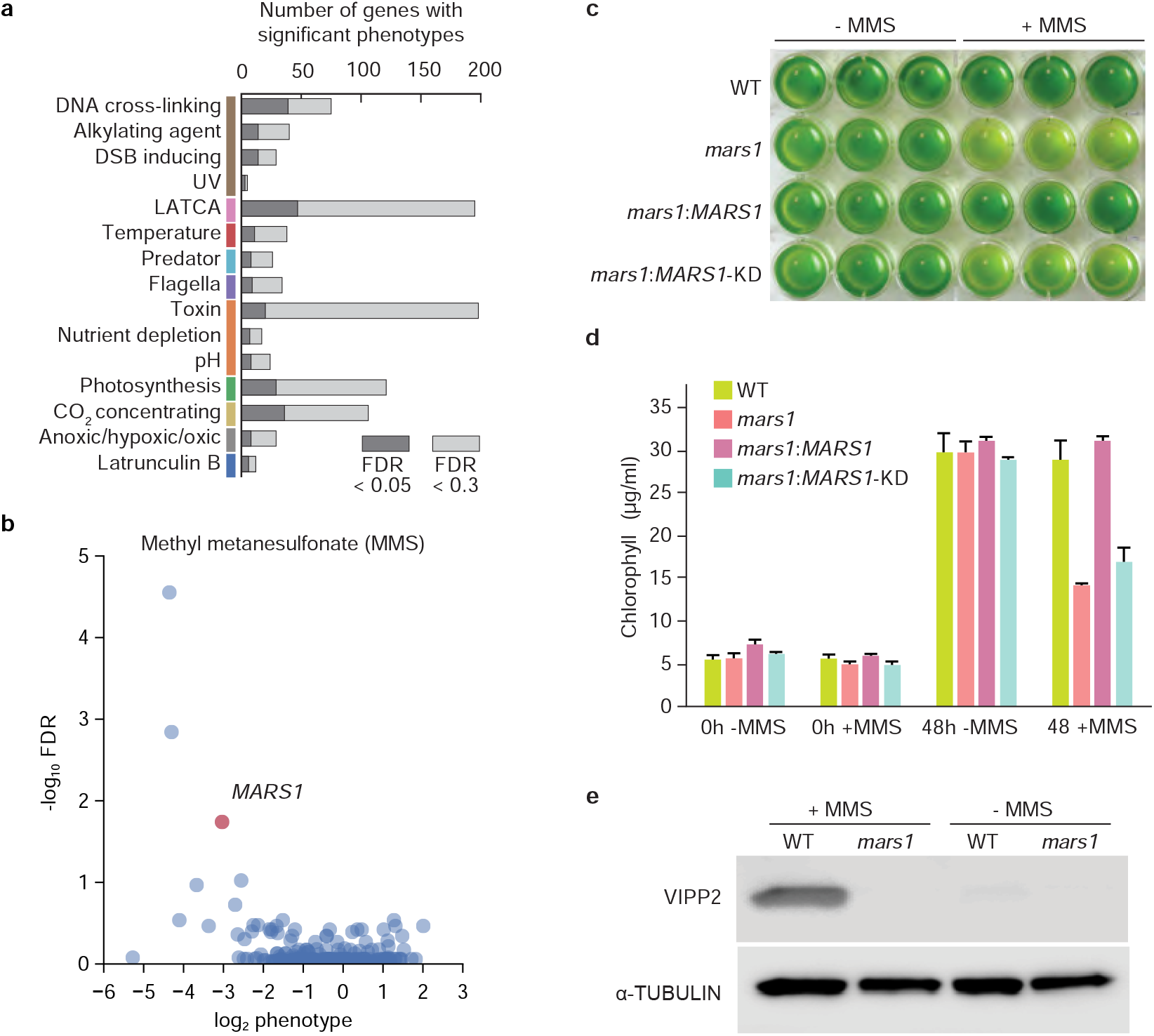
Multiple alleles provide high confidence and reveal novel phenotypes. **a**. The number of genes with significant phenotypes in each class of screen is shown for two false discovery rate (FDR) thresholds. **b**. FDR is plotted against log2 median phenotype for all genes in the methyl methanesulfonate (MMS) screen. **c**. Growth assay of wild-type (WT), *mars1*, *mars1:MARS1,* and *mars1:MARS1 KD* cells after 48 h in the presence or absence of MMS. Three biological replicates were used for each strain. For more details, see materials and methods. **d**. Average chlorophyll concentrations of the liquid cultures shown in Figure 3c. **e**. Immunoblot analysis of VESICLE-INDUCING PROTEIN IN PLASTIDS (VIPP2), a downstream target of MARS1, in WT and *mars1* cells in the presence or absence of MMS.

As an example of how individual gene-phenotype relationships advance our understanding, we made the unexpected observation that mutants in the gene encoding the chloroplast unfolded protein response (cpUPR) kinase, MUTANT AFFECTED IN CHLOROPLAST-TO-NUCLEUS RETROGRADE SIGNALING (MARS1)^17^, were sensitive (FDR <10^−9^) to the DNA damaging agent methyl methanesulfonate (MMS) (Fig. 3b). We validated this phenotype in a separate growth assay and showed that the MMS sensitivity of these mutants is rescued by complementation with a wild-type copy of *MARS1* but not by a kinase-dead version (Fig. 3c-d). We also determined that treatment with MMS leads to induction of VESICLE-INDUCING PROTEIN IN PLASTIDS 2 (VIPP2), a highly selective cpUPR marker, in wild-type cells but not in mutants lacking *MARS1* (Fig. 3e). These results illustrate the value of our high-throughput data and suggest the intriguing possibility that the cpUPR is activated via MARS1 upon DNA damage or protein alkylation and has a protective role against these stressors.

## From phenotypes to pathways

To facilitate data visualization and to help the prediction of functions for poorly-characterized genes in our dataset, we used the principle that genes whose mutants have similar phenotypes tend to function in the same pathway^5^. We clustered the 684 genes with high-confidence phenotypes based on the similarity of their phenotypes across different treatments (Fig. 4a and Extended Data File 2). The correlation of phenotypes was largely unrelated to transcriptional expression correlation, suggesting that the two approaches provide complementary information (Extended Data Fig. 3, Table S8). We named some of our gene clusters based on the presence of previously characterized genes or based on the conditions that produced the most dramatic phenotypes in a cluster (Fig. 4b-g). Below, we provide examples of how the data recapitulate known genetic relationships and provide insights into the functions of poorly characterized genes.

**Figure 4.**
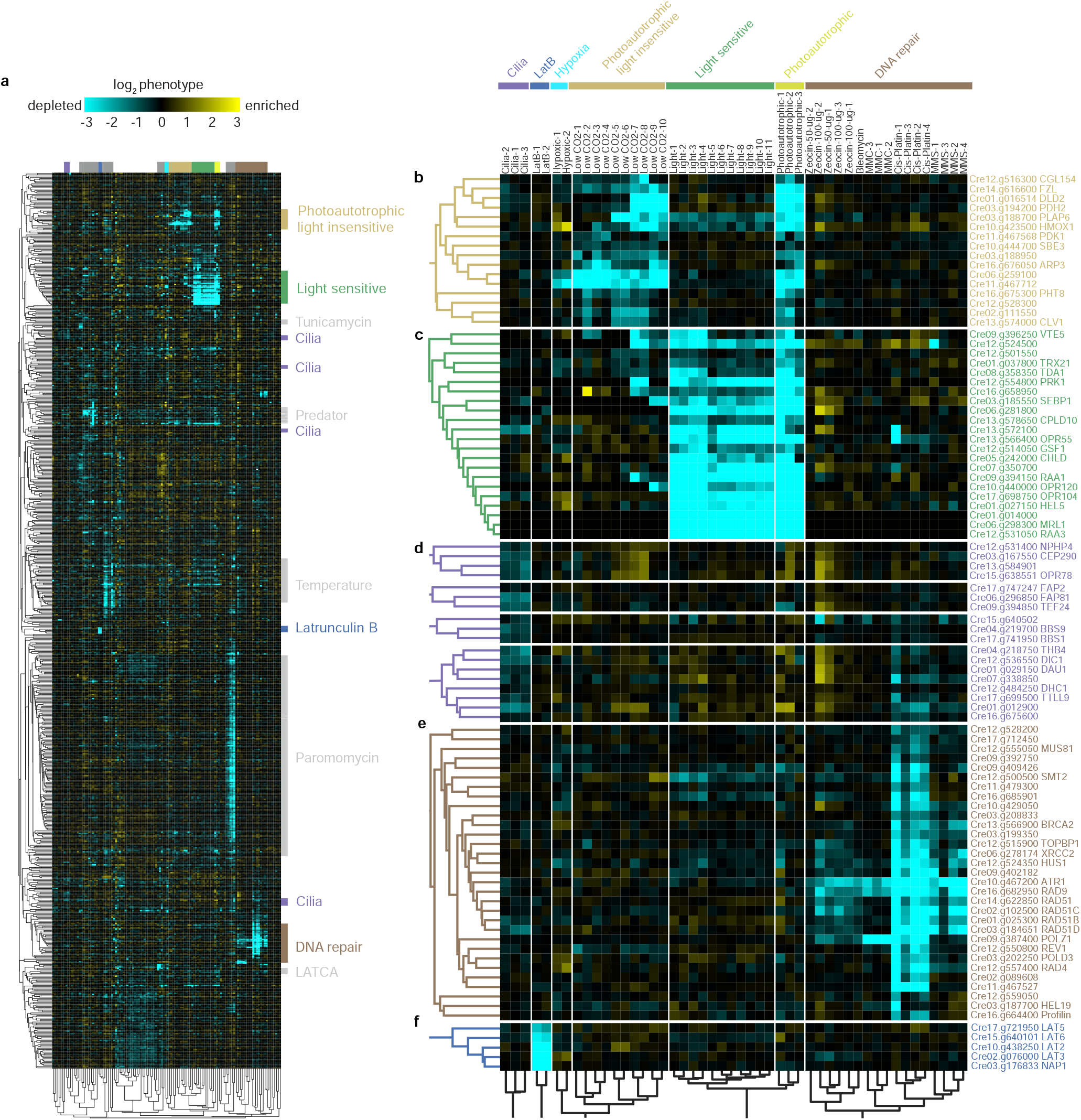
Similarity of mutant phenotypes places genes into pathways and reveals novel players. **a**. 684 genes were clustered based on the similarity of their phenotypes across 120 screens. b-f. Examples of how sub-clusters enriched in specific pathways predict novel genes in these pathways: b, non-photoautotrophic light-insensitive; c, non-photoautotrophic light-sensitive; d, cilia; e, DNA damage-sensitive; f, Latrunculin B-sensitive.

## Essential DNA repair pathways are conserved to green algae

DNA damage repair pathways are among the best-characterized and most highly conserved across all organisms^19,20^, thus, they serve as a useful test case of the quality of our data. In our dataset, 21 homologues of known DNA repair proteins formed a large cluster (Fig. 4e), demonstrating the quality of our phenotypic data, validating our ability to identify that these genes work in a common pathway, and extending the conservation of their functions to green algae.

Mutants for various DNA repair genes exhibit expected differences in their sensitivities to different types of DNA damage: 1) DNA double-strand breaks (zeocin and bleomycin), 2) DNA crosslinks (Mitomycin C and cisplatin), and 3) DNA alkylation (MMS). For example, mutants exhibiting sensitivity to all DNA damage conditions included cells lacking upstream DNA damage-sensing kinase ATAXIA TELANGIECTASIA AND RAD3-related protein (ATR, encoded by Cre10.g467200)^21^; as well as mutants lacking the cell cycle checkpoint control protein RADIATION SENSITIVE 9 (RAD9, encoded by Cre16.g682950) or its binding partner HYDROXYUREA-SENSITIVE 1 (HUS1, encoded by Cre12.g524350)^22^. Mutants specifically sensitive to the double-strand break-inducing agents zeocin and bleomycin included the upstream sensor of double-strand breaks, the kinase ATAXIA-TELANGIECTASIA MUTATED (ATM, encoded by Cre13.g564350)^23^ (Table S6); as well as DNA POLYMERASE THETA (POLQ, encoded by Cre16.g664301), which facilitates error-prone double-strand break repair and can maintain genome integrity when other repair pathways are insufficient^24,25^ (Table S6). Mutants specifically sensitive to the DNA crosslinker cisplatin included cells with genetic lesions in the helicases REGULATOR OF TELOMERE ELONGATION HELICASE 1 (RTEL1, encoded by Cre02.g089608)^26^, in FANCONI ANEMIA COMPLEMENTATION GROUP M (FANCM, encoded by Cre03.g208833), and in the crossover junction endonuclease METHANSULFONATE UV SENSITIVE 81 (MUS81, encoded by Cre12.g555050).

Our data suggest several instances where a given factor is required for the repair of a specific kind of DNA damage in Chlamydomonas but not in Arabidopsis, or vice-versa, suggesting lineage-specific differences in how DNA damage is repaired. For example, Chlamydomonas *fancm* mutants are sensitive to the DNA crosslinker cisplatin, while Arabidopsis *fancm* mutants are not^27^. Conversely, Arabidopsis *mus81* mutants are sensitive to the alkylating agent MMS and the DNA crosslinker Mitomycin C^28^, while Chlamydomonas *mus81* mutants were not.

Taken together, our data suggest that the core eukaryotic DNA repair machinery defined in other systems is generally conserved in green algae. Moreover, the observation of expected phenotypes illustrates the quality of the presented data and the utility of the platform for chemical genomic studies.

## The data allow classification of photosynthesis genes

Our data allowed the classification of 38 genes whose disruption leads to a photoautotrophic growth defect^12^ into two clusters. One cluster consisted of genes whose disruption confers sensitivity to light when grown on medium supplemented with acetate while the other contained genes whose disruption does not (Fig. 4b-c, Extended Data File 2).

The light-sensitive cluster (Fig. 4c) included genes encoding core photosynthesis components and biogenesis factors such as the pmRNA trans-splicing factors *RNA MATURATION Of PSAA* (*RAA1*)^29^, *RAA3*^30^, *OCTOTRICOPEPTIDE REPEAT 120* (*OPR12*0), and *OPR104*^31^; Photosystem II biogenesis factor *CONSERVED IN PLANT LINEAGE AND DIATOMS 10* (*CPLD10*)^*31,32*^ the chlorophyll biogenesis factor *Mg-CHELATASE SUBUNIT D* (*CHLD*)^33^; the ATP synthase translation factor *TRANSLATION DEFICINET ATPase 1* (*TDA1*)^34^; the Rubisco mRNA stabilization factor *MATURATION OF RBCL 1* (*MRL1*)^35^; and the Calvin-Benson-Bassham cycle enzymes *SEDOHEPTULOSE-BISPHOSPHATASE 1* (*SEBP1*)^36^ and *PHOSPHORIBULOKINASE 1* (*PRK1*)^37^. Several highly conserved but poorly characterized genes are also found in this cluster, including the putative Rubisco methyltransferase Cre12.g524500^38^, the putative thioredoxins Cre01.g037800, Cre06.g281800, and Cre13.g572100; as well as four *Chlorophyta*-specific genes. The mutant phenotypes of these poorly-characterized genes and their presence in this light-sensitive cluster together suggest that their products could mediate the biogenesis, function, or regulation of core components of the photosynthetic machinery.

The light-insensitive cluster (Fig. 4b) contained known and novel components of the algal CO_2_-concentrating mechanism (CCM), as detailed below.

## Novel CO_2_-concentrating mechanism components

The CO_2_-concentrating mechanism increases the CO_2_ concentration around the CO_2_ fixing enzyme Rubisco, thus enhancing the rate of carbon uptake. The mechanism uses carbonic anhydrases in the chloroplast stroma to convert CO_2_ to HCO_3_^−^, which is transported into the lumen of the thylakoid membranes that traverse a Rubisco-containing structure called the pyrenoid^39^. There, the lower pH drives the conversion of HCO_3_^−^ back into concentrated CO_2_ that feeds Rubisco^39^. Mutants deficient in the CO_2_-concentrating mechanism are unable to grow photoautotrophically in air, but their photoautotrophic growth is rescued in 3% CO_2_39. We observed this phenotype for one or more alleles of genes whose disruption was previously shown to disrupt the CCM (Table S4), including genes encoding the chloroplast envelope HCO_3_^−^ transporter LOW CO_2_ INDUCIBLE GENE A (LCIA)^40^, and the thylakoid lumen CARBONIC ANHYDRASE 3 (CAH3)^41^, the stromal carbonic anhydrase LOW CO_2_ INDUCIBLE GENE B (LCIB)^42^, the master transcriptional regulator CCM1/CIA5^43,44^, and the pyrenoid structural protein STARCH GRANULES ABNORMAL 1 (SAGA1)^45^(Table S6).

Similarly, we observed high CO_2_ rescue of photoautotrophic growth defects for mutants in multiple poorly characterized genes in the light-insensitive cluster, suggesting that many of these genes are novel components in the CO_2_-concentrating mechanism. These genes formed a cluster with *SAGA1*^45^, the only previously known CO_2_-concentrating mechanism gene with enough alleles to be present in the cluster. We named one of these components, Cre06.g259100, *SAGA3* because its protein product shows homology to the two pyrenoid structural proteins SAGA1 and SAGA2^46^ (Extended Data Fig. 4). Consistent with a role in the CO_2_-concentrating mechanism, SAGA3 localizes to the pyrenoid^47^. We also observed this phenotype in mutants lacking the pyrenoid starch sheath-localized *STARCH BRANCHING ENZYME 3* (*SBE3*)^48^, suggesting that this enzyme plays a key role in the biogenesis of the pyrenoid starch sheath, a structure surrounding the pyrenoid that has recently been shown to be important for pyrenoid function under some conditions^49^. Our cluster also contains FUZZY ONIONS (FZO)-like (FZL), a dynamin-related membrane remodeling protein involved in thylakoid fusion and light stress; mutants in this gene have pyrenoid shape defects^50^. Our results suggest that thylakoid organization influences pyrenoid function. Additional genes showing similar phenotypes included *CLV1* (encoded by Cre13.g574000), a predicted voltage-gated chloride channel that we hypothesize is important for regulating the ion balance in support of the CO_2_-concentrating mechanism, or alternatively, may directly mediate HCO3-transport; a protein containing a Rubisco-binding motif (encoded by Cre12.g528300); and a predicted Ser-Thr kinase (Cre02.g111550). The kinase is a promising candidate for a regulator in the CO_2_-concentrating mechanism, as multiple CCM components are known to be phosphorylated^51–53^, but no kinase had previously been shown to have a CCM phenotype.

Also in this cluster of high CO_2_ rescue genes are the predicted *PYRUVATE DEHYDROGENASE 2* (*PDH2)* (Cre03.g194200) and the predicted DIHYDROLIPOYL DEHYDROGENASE (*DLD2)* (Cre01.g016514). We hypothesize that these proteins are part of a glycine decarboxylase complex that functions in photorespiration, a pathway that recovers carbon from the products of the Rubisco oxygenation reaction. *PDH2* was found in the pyrenoid proteome^54^, suggesting the intriguing possibility that glycine decarboxylation may be localized to the pyrenoid, where the recovered CO_2_ could be captured by Rubisco.

## Novel genes with roles in cilia function

Chlamydomonas cells swim using two motile cilia. To identify mutants with abnormal cilia function, we separated mutants based on swimming ability by placing the pool of mutants in a vertical column and collecting the supernatant and pellet. In this assay, mutants with altered swimming behavior were enriched in GO terms such as “dynein complex,” which comprises motor proteins involved in ciliary motility (Fig. 2e). 18 genes were represented by enough alleles to provide high confidence (FDR < 0.3) that their disruption produces a defect in swimming (Fig. 4d). These genes were enriched (*P*=0.0075, Fisher’s exact test) in genes encoding proteins found in the Chlamydomonas flagella proteome^55^. Half of these genes or their orthologs have previously been associated with a cilia-related phenotype in Chlamydomonas and/or mice (Table S9).

In our analysis, these 18 genes formed four clusters that appeared to sub-classify their function (Fig. 4d). The first cluster is enriched in known regulators of ciliary membrane composition and includes *NEPHROCYSTIN-4-LIKE PROTEIN (NPHP4*)^56^; its physical interactor *TRANSMEMBRANE PROTEIN 67* (*TMEM67*, also named *MECKEL SYNDROME TYPE 3* [MKS3] in mammals), which has been implicated in photoreceptor intraciliary transport^57^; and *CENTRIOLE PROTEOME PROTEIN* 290 (*CEP290*)^58^. We validated the swimming defect of *tmem67* and observed that the mutant has shorter cilia (Extended Data Fig. 5). The poorly-annotated gene Cre15.g638551 clusters with these genes, suggesting that it may also regulate ciliary membrane composition.

The second cluster contains *BARDET-BIEDL SYNDROME 1 PROTEIN 1 (BBS1*) and *BBS9*, components of the Bardet-Biedl syndrome-associated complex that regulates targeting of proteins to cilia^59^. The poorly annotated gene Cre15.g640502 clustered with these genes, suggesting that it may also play a role in targeting proteins to cilia.

The third cluster contains eight genes, four of which relate to the dynein complex. These genes include the ciliary dynein assembly factor *DYNEIN ASSEMBLY LEUCINE-RICH REPEAT PROTEIN (DAU1*)^60,61^; *OUTER DYNEIN ARM* (*ODA*); *DYNEIN ARM INTERMEDIATE CHAIN* 1 (*DIC1*)^62^; *DYNEIN HEAVY CHAIN* 1 (*DHC1*)^63^; and *TUBULIN-TYROSINE LIGASE* 9, (*TTLL9*), which modulates ciliary beating through the addition of a polyglutamate chains to alpha tubulin^64^. The predicted thioredoxin peroxidase gene Cre04.g218750 and three poorly annotated genes (Cre07.g338850, Cre01.g012900, and Cre16.g675600) clustered with these genes, suggesting possible roles in dynein assembly or regulation.

The fourth cluster contains three poorly characterized genes, *FLAGELLA ASSOCIATED PROTEIN2* (*FAP2*), *FLAGELLA ASSICIATED PROTEIN 81* (*FAP81)*, and *TEF24*. The protein encoded by FAP81 (Cre06.g296850) was identified in the Chlamydomonas cilia proteome^55^, and its human homolog DELETED IN LUNG AND ESOPHAGEAL CANCER PROTEIN 1 (*DLEC1*) localizes to motile cilia^65^. We validated the swimming defect of the *fap81* mutant and establisheds that it has shorter cilia (Extended Data Fig. 5). The localization to motile cilia in humans and our finding that mutating the encoding gene leads to a ciliary motility defect together suggests the intriguing possibility that impaired cilia motility contributes to certain lung and esophageal cancers.

## Novel genes required for actin cytoskeleton integrity

Our analysis revealed a group of genes whose mutation render cells sensitive to LatB (Fig. 4g). LatB binds to monomers of actin, one of the most abundant and conserved proteins in eukaryotic cells, and prevents actin polymerization^66^ (Fig. 5a). LatB was first discovered as a small molecule that protects the sea sponge *Latrunculina magnifica* from predation by fish^67^, and is an example of the chemical warfare that organisms use to defend themselves and compete in nature (Fig. 5b).

**Figure 5:**
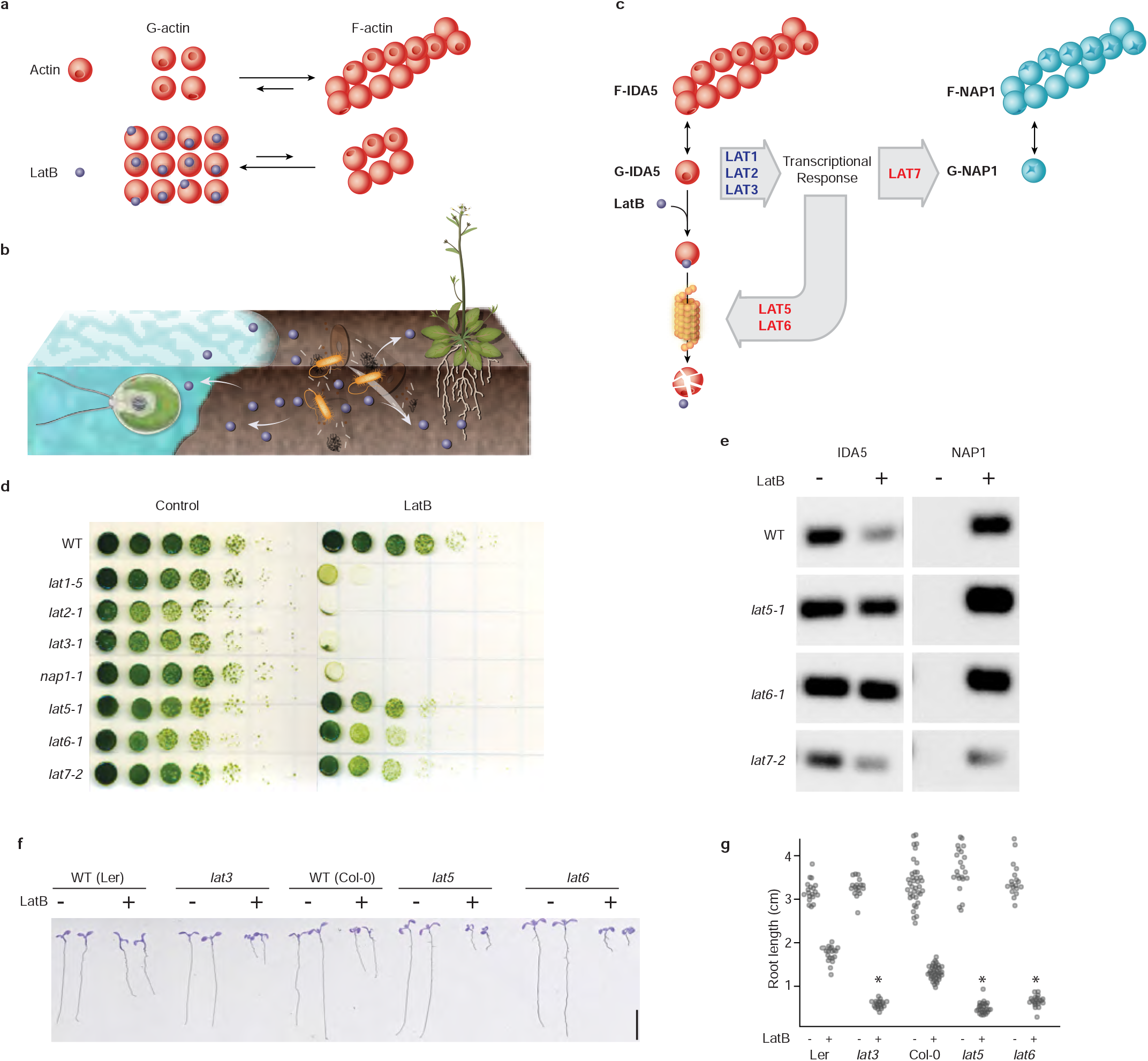
The approach revealed novel conserved components of a defense mechanism against cytoskeleton inhibitors. **a**. Latrunculin B (LatB) interferes with actin polymerization. **b**. Soil microorganisms deploy actin inhibitors for a competitive advantage in their environment. **c**. Chlamydomonas responds to actin inhibition by degrading its conventional actin IDA5 and upregulating an alternative actin, NAP1. **d**. Growth of new *lat* mutants identified in this study (*lat5-1*, *lat6-1*, and *lat7-2*) was compared with previously isolated *lat1-5*, *lat2-1*, *lat3-1*, and *nap1-1* mutants^68^ in the absence (control) and presence (LatB) of 3 μM LatB. **e**. Immunoblot of conventional (IDA5) and alternative (NAP1) actins shows that *lat5-1* and *lat6-1* are deficient in actin degradation, while *lat7-2* lacks proper induction of the non-canonical actin (NAP1) when exposed to an actin inhibitor. **f**. The F-actin homeostasis pathway is conserved between green algae and plants. Mutants in Arabidopsis genes homologous to Chlamydomonas *lat3*, *lat5,* and *lat6* show sensitivity to LatB as decreased root length. **g**. Quantification of root length in Arabidopsis mutants. Asterisks mark significant changes based on two-way ANOVA, p <0.05.

Chlamydomonas protects itself against LatB-mediated inhibition of its conventional actin INNER DYNEIN ARM5 (IDA5) by upregulating the highly divergent actin homologue NOVEL ACTIN-LIKE PROTEIN1 (NAP1), which appears to perform most of the same functions as actin but is resistant to inhibition by LatB^68^. Upon inhibition of IDA5 by LatB, IDA5 is degraded and divergent actin NAP1 is expressed^68^. The expression of *NAP1* is dependent on three other known genes, *LatB-SENSITIVE* (*LAT1-LAT3)* (Fig. 5c); thus, mutants lacking any of these four genes are highly sensitive to LatB^68^.

Our phenotype data revealed three novel components of this F-actin homeostasis pathway, which we named LAT5 (Cre17.g721950), LAT6 (Cre15.g640101) and LAT7 (Cre11.g482750). LAT5 and LAT6 clustered together with three previously known components of the pathway: *NAP1*, *LAT2* and *LAT3*; and disruption of all six genes rendered cells sensitive to LatB (Table S6). Mutants in all three novel components show a relatively mild phenotype when compared with those mutants in *LAT1-LAT3* (Fig. 5d), illustrating the sensitivity of our phenotyping platform.

Ubiquitin proteosome-mediated proteolysis of IDA5 has been hypothesized to drive the degradation of IDA5 and promote the formation of F-NAP1^69^, but the factors involved were unknown. *LAT5* and *LAT6* encode predicted subunits of a SKP1, CDC53/CULLIN, F-BOX RECEPTOR (SCF) E3 ubiquitin ligase, whose homologs promote the degradation of target proteins^70^ the disruption of *LAT5* and *LAT6* impaired degradation of IDA5 upon LatB treatment, suggesting that LAT5 and LAT6 mediate IDA5 degradation (Fig. 5e). *LAT7* encodes a predicted importin, and its disruption impairs NAP1 accumulation after LatB treatment (Fig. 5e), suggesting that nuclear import is required for NAP1 biosynthesis.

It was previously not clear how broadly conserved this F-actin homeostasis pathway is. We found that the land plant model Arabidopsis has homologs of IDA5, NAP1, LAT3, LAT5, LAT6 and LAT7. We observed that Arabidopsis mutants disrupted in *LAT3*, *LAT5,* and *LAT6* are sensitive to LatB treatment (Fig. 5f-g), which was not expected *a priori*, suggesting that this pathway for actin cytoskeleton integrity and the gene functions identified here are conserved in land plants.

## Discussion

In this work, we determined the phenotypes of 58,101 Chlamydomonas mutants across a broad variety of growth conditions. We observed a phenotype for mutants representing 13,840 genes, providing a valuable starting point for characterizing the functions of thousands of genes. Mutant phenotypes are searchable at chlamylibrary.org, and individual mutants can be ordered from the Chlamydomonas Resource Center.

We provided several examples of how the data enable discovery of novel gene functions and phenotypes in algae and plants. We validated our discovery of three novel genes in the actin cytoskeleton integrity pathway, obtained insights into their molecular functions, and found that this pathway appears to be conserved in land plants. We validated our discovery of cilia function defects for two novel genes and our observation of an unexpected sensitivity of the chloroplast unfolded protein response to the alkylating agent MMS. We also discussed how our data provide insights and candidate genes in other pathways including DNA damage repair, photosynthesis, and the CO_2_ concentrating mechanism.

This work illustrates the value of using a microbial photosynthetic organism for discovering novel gene functions on a large scale. We hope that the genotype-phenotype relationships identified here will guide the characterization of thousands of genes, with potential applications in agriculture, the global carbon cycle, and our basic understanding of cell biology.

## Acknowledgments

We thank Matthew Cahn for developing and improving the CLiP website; Xuhuai Ji at the Stanford Functional Genomics Facility and Ziming Weng at the Stanford Center for Genomics and Personalized Medicine for deep sequencing services; Kathryn Barton, Winslow Briggs, and Zhi-Yong Wang for providing lab space; Ted Raab for providing lab equipment; Jacob Robertson and Nina Ivanova for helping with library maintenance; Members of the Dinneny, Jonikas, and Jinkerson Labs, Tingting Xiang, and Alexandra Wilson for constructive suggestions on the manuscript.

This project was supported by grants from the National Institutes of Health (DP2-GM-119137), the National Science Foundation (MCB-1914989 and MCB-1146621), and the Simons Foundation and Howard Hughes Medical Institute (55108535) awarded to M.C.J.; a German Academic Exchange Service (DAAD) research fellowship awarded to F.F.; Simons Foundation fellowships of the Life Sciences Research Foundation awarded to R.E.J. and J.V.-B.; EMBO long term fellowship (ALTF 1450-2014 and ALTF 563-2013) awarded to J.V.-B and S.R.; an NSF MCB Grant (1818383) awarded to M.O.; and a Swiss National Science Foundation Advanced PostDoc Mobility Fellowship (P2GEP3_148531) awarded to S.R; a Simons Foundation and Howard Hughes Medical Institute (55108515) Faculty Scholars grant awarded to J.R.D. Work in the Merchant laboratory is supported by a cooperative agreement with the US Department of Energy Office of Science, Office of Biological and Environmental Research program under Award DE-FC02-02ER63421.

## Author Contributions

S.W. and K.K.N. performed rose bengal treatments; R.G.K., Y.K., and A.R.G. performed anoxia and high light treatments; J.V.-B., F.F., and R.E.J. prepared mutant pools, performed all other treatments, and processed all samples; S.C. provided the LATCA compounds; M.M., J.O., C.P., and M.N. validated the active LATCA compounds; M.M. performed chemoinformatics analysis; M.O. validated LatB phenotypes in Chlamydomonas and performed immunoblots; S.R. and P.W. validated the *mars1* phenotype; W.P. guided statistical analysis and website development; P.A.S. and S.M. provided transcriptomics data; X.L. provided early access to the Chlamydomonas mutant library; J.V.-B. and J.R.D. confirmed Arabidopsis phenotypes; J.V.-B., F.F., M.C.J., and R.E.J. designed experiments, analyzed, and interpreted the data; J.V.-B., F.F., M.C.J., and R.E.J. wrote the manuscript with input from all authors.

The authors declare no competing financial interests. J.V.-B., F.F., M.C.J., and R.E.J. wish to note that a provisional patent application on aspects of the findings has been submitted.

## Additional Information

Supplementary information is available for this paper. J.V.B and F.F. contributed equally to this work. Correspondence and requests for materials should be addressed to J.D., M.C.J. and R.E.J.

---

**Extended Data Figure 1.**
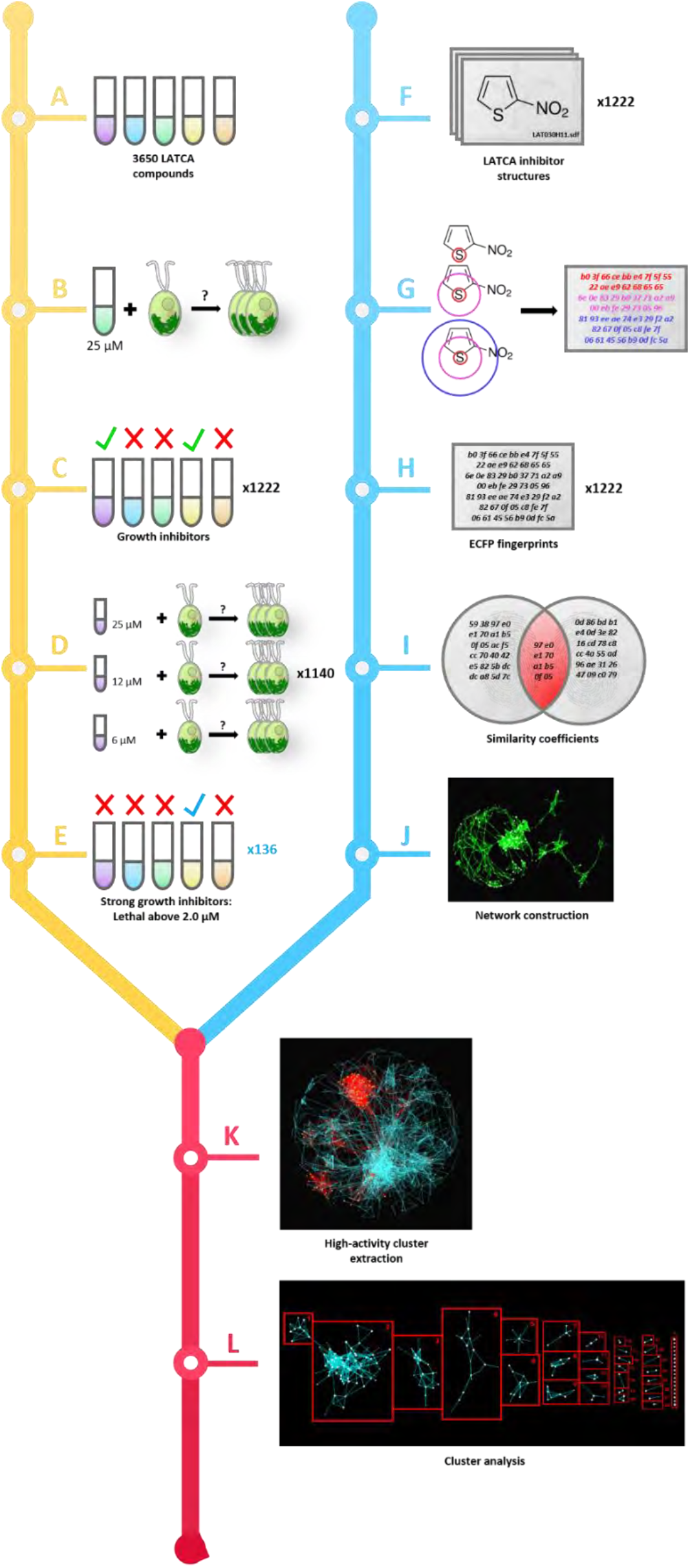
A screen of the chemical library “LATCA” identified 1,222 inhibitors of Chlamydomonas growth, 136 of which are active at 2 μM or less. Phase 1 of the LATCA screen is depicted in A-E: The growth rate of wild-type Chlamydomonas (cMJ030) was evaluated in TAP and TP in the presence of 3,650 LATCA compounds. 1,222 out of the 3,650 LATCA compounds reduced growth by 90% or more at 25 μM (A–C). Dose-response experiments were performed in TAP media with 1,140 out of the 1,222 highly active compounds that reduced growth at 2 μM or less (D,E). Phase 2 of the LATCA screen is depicted in F–J: Structural data files (SDFs) were acquired for all LATCA inhibitors (F) and converted into numerical fingerprints (extended-connectivity fingerprints; ECFPs) (G, H). ECFPs were then used to compute the structural similarity of pairs of compounds using Tanimoto coefficients (I). The set of Tanimoto coefficients between all pairs of inhibitors was condensed into a usable network (J). Phase 3 of the LATCA screen is depicted in K and L: Data from A–E was used to further reduce the similarity network from J to 28 clusters of structures exhibiting high levels of growth inhibition along with a group of singleton structures (*) that did not cluster. Table S3 summarizes data A–E and shows cluster annotations from L; see also Extended Data File 1 for all chemical structures from L.

**Extended Data Figure 2.**
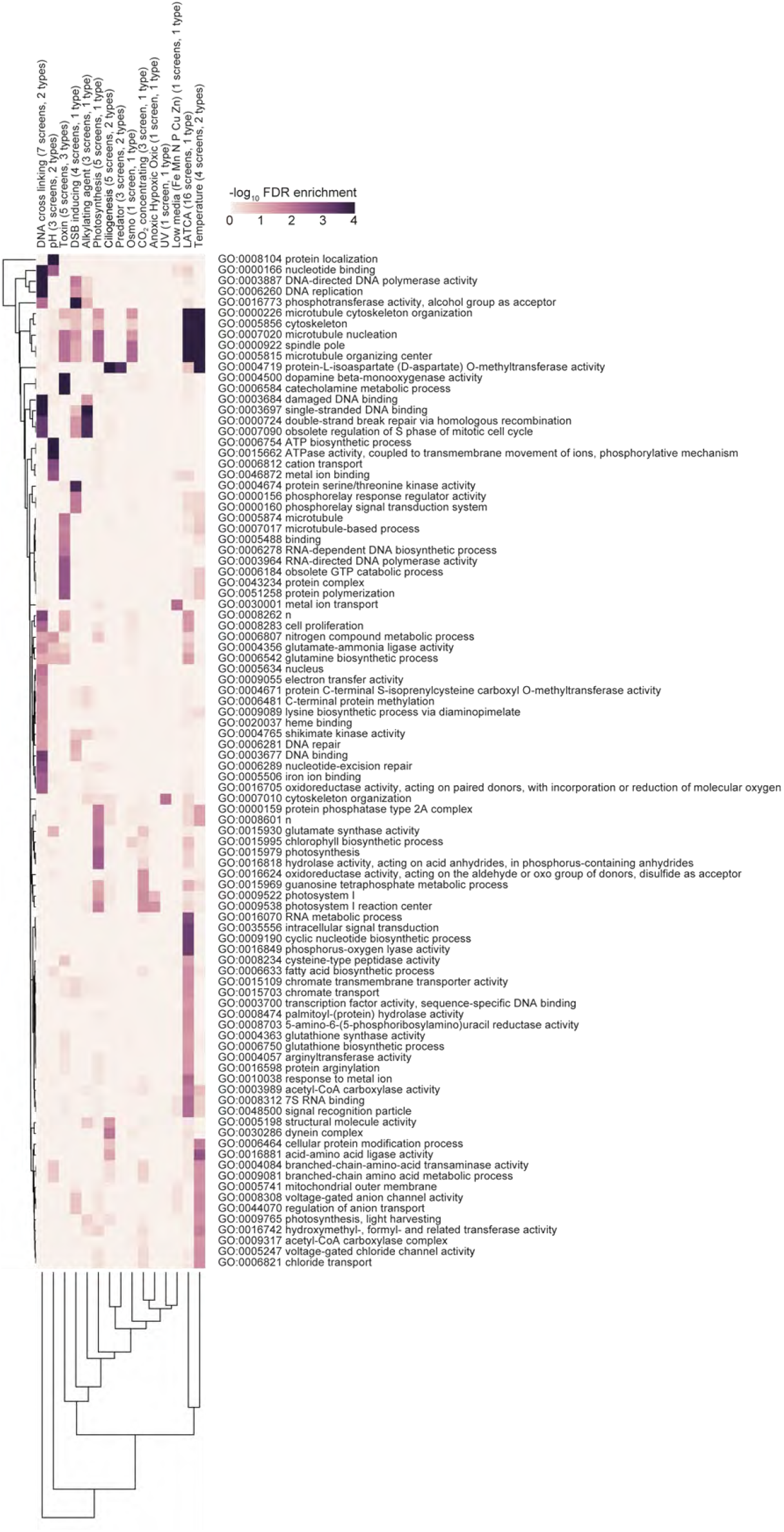
Full GO term enrichments. Gene Ontology term analysis reveals enrichment of biological functions observed for specific screens. GO; FDR<0.05.

**Extended Data Figure 3.**
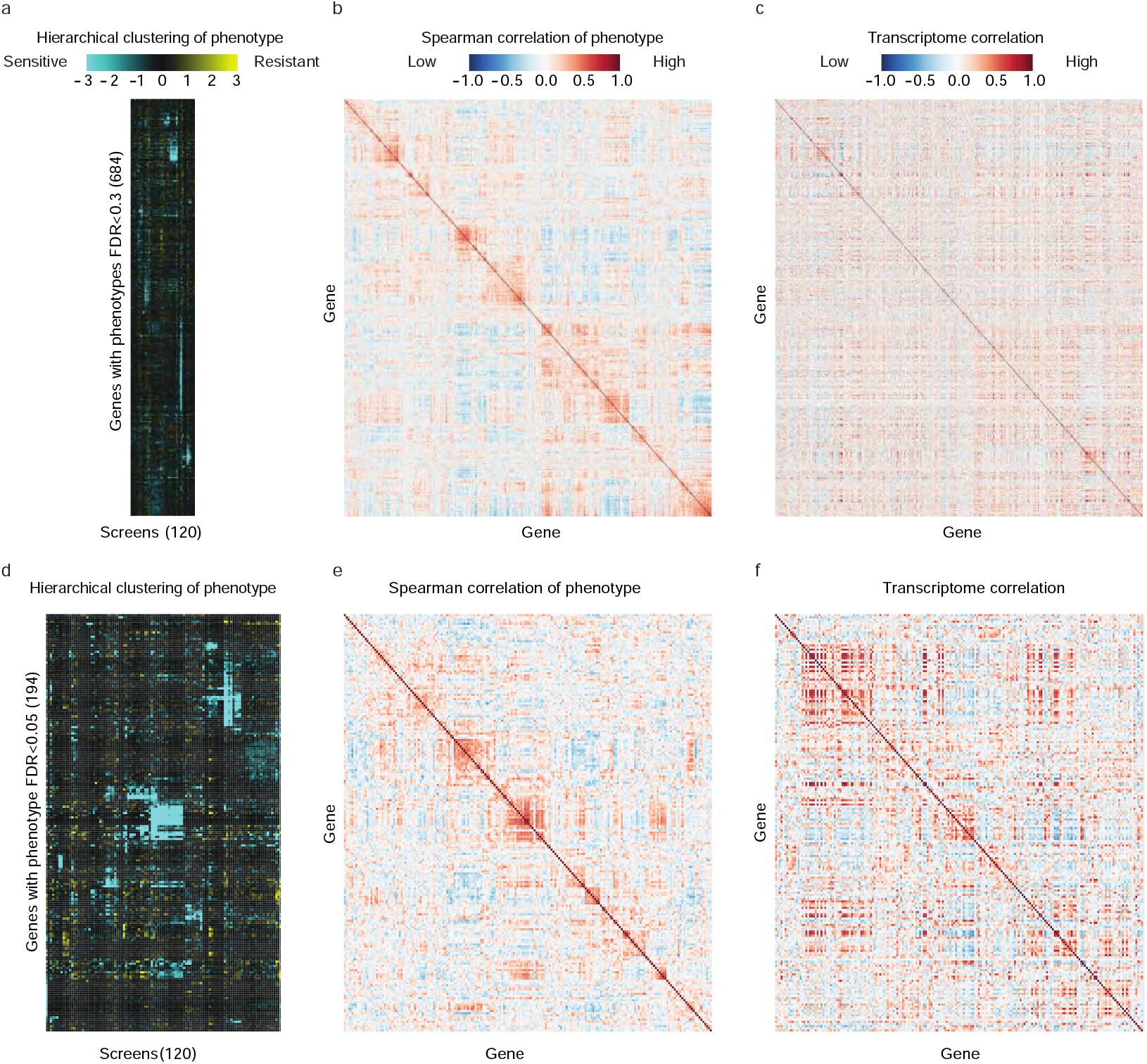
Comparison of phenotypic and transcriptomic correlations. **a**. 684 genes were clustered based on the similarity of their phenotypes across 120 screens. **b**. Spearman correlation matrix of phenotypes (FDR <0.3). **c**. Transcriptome correlation of gene with phenotype (FDR <0.3). **d**. 194 genes were clustered based on the similarity of their phenotype across 120 screens. **e**. Spearman correlation matrix of phenotypes (FDR <0.05). **f**. Transcriptome correlation of gene with phenotype (FDR <0.05). Data can be found in Table S8.

**Extended Data Figure 4.**
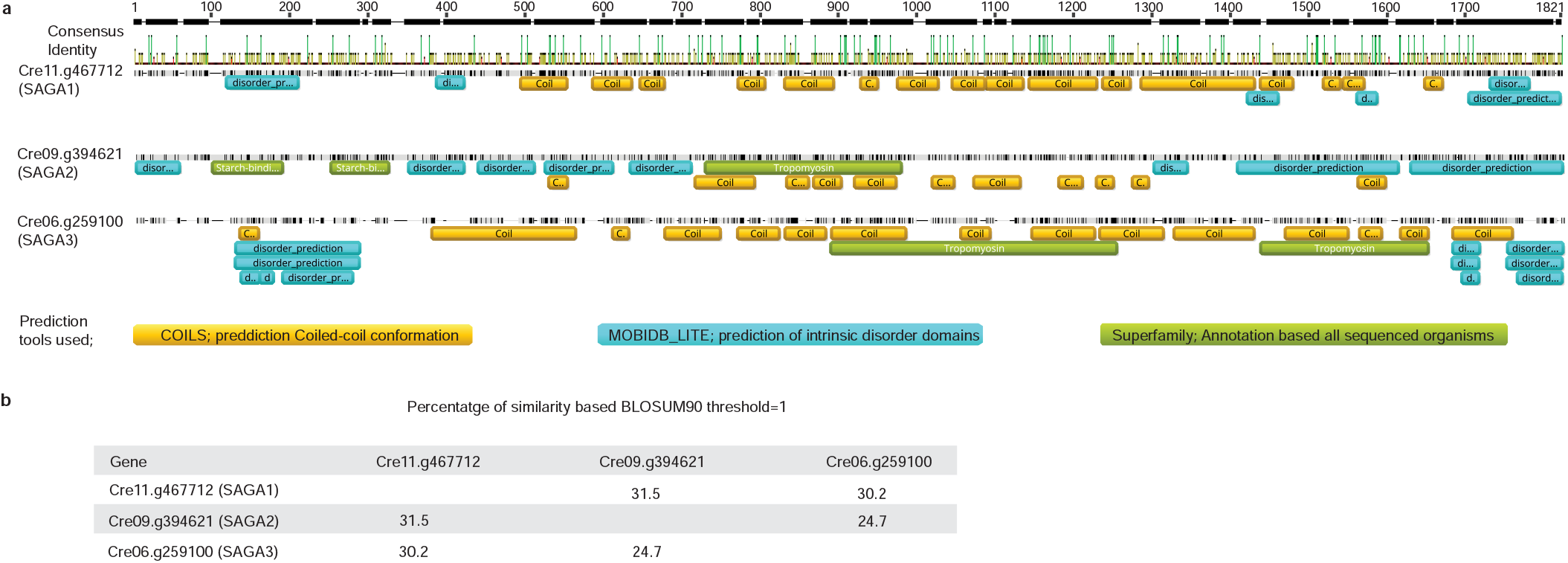
SAGA protein alignments. **a**. Alignments of SAGA1, SAGA2, and SAGA3. Domain annotation was based on three different tools under the *Geneious* visualization platform. **b**. BLOSUM90 alignments between SAGA proteins.

**Extended Data Figure 5.**
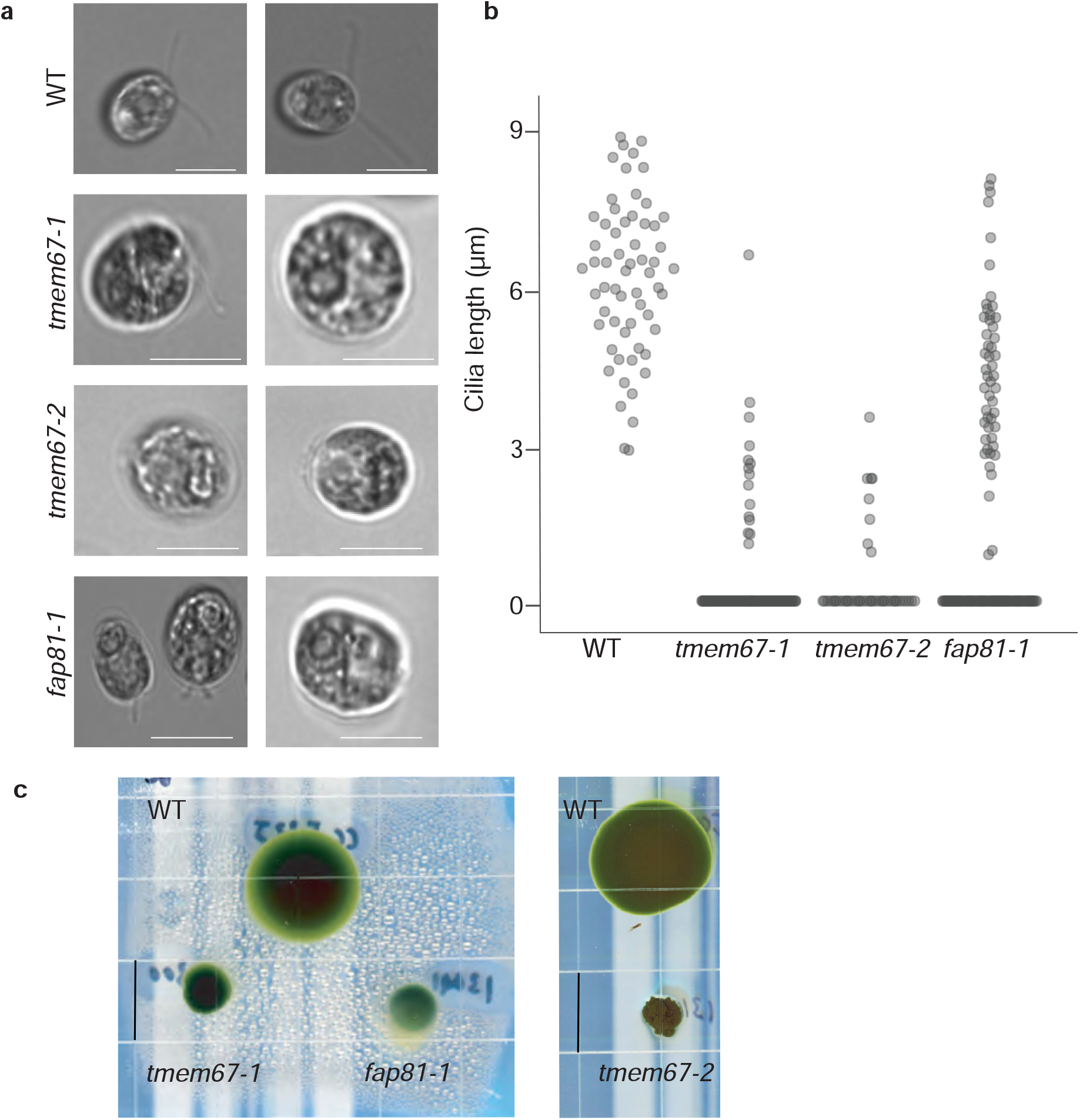
Validation of cilia mutant phenotypes. **a**. Bright-field microscopy microscope images of cilia mutants show defects in ciliary length. Scale bar: 10 μm. **b**. Quantification of cilia length. **c**. Swimming behavior of mutants, as determined by growth on TAP medium solidified with 0.15% agar. Scale bar: 5cm.

## LIST OF SUPPLEMENTARY MATERIALS

Supplementary materials can be downloaded from the Chlamylibrary website: https://www.chlamylibrary.org/download

Table S1 | Source material used for screens

Table S2 | List of screens

Table S3 | Initial LATCA screen and validation; cluster groups

Table S4 | Raw mutant phenotypes across all conditions

Table S5 | FDRs for GO term enrichment

Table S6 | FDRs for all genes by all screens

Table S7 | High-confidence gene-phenotype relationships

Table S8 | Phenotypic and transcriptomic correlations of genes with high-confidence phenotypes

Table S9 | Cluster annotations with yeast, mouse, and Arabidopsis orthologs

Table S10 | Primers used in this study

Table S11 | Raw and normalized read counts

Table S12 | List of samples that were averaged

Table S13 | Chlamydomonas and Arabidopsis strains used in this study

Extended Data File 1 | LATCA compound structures

Extended Data File 2 | Java TreeView files of FDR less than 0.3 gene clusters

## METHODS

### Library maintenance

The Chlamydomonas mutant collection^12^ was maintained by robotically passaging 384-colony arrays to fresh medium using a Singer RoToR robot (Singer Instruments, 704 Somerset, UK). The mutant collection was grown on 1.5% agar Tris-Acetate-Phosphate (TAP) medium with modified trace elements^71^ in complete darkness at room temperature. The routine passaging interval of four weeks for library maintenance was shortened to two weeks during the time period of pooled screens to increase cell viability.

### Screening of the Library of AcTive Compounds in Arabidopsis (LATCA) to identify Chlamydomonas growth inhibitorss

The Library of AcTive Compounds in Arabidopsis (LATCA)^13^ was used to identify molecules capable of inhibiting growth in wild-type Chlamydomonas (cMJ030). We found that 1,222 of these 3,650 LATCA compounds reduce growth by 90% at 25 μM (Table S3). Due to resource limitations, we could not perform competitive growth experiments with all 1,222 active chemicals. Hence, we further selected the most active compounds and analyzed their structural similarity to identify the most diverse set of compounds for the final screen. We performed dose-response experiments with 1,140 compounds, validated activity for 954 compounds, and identified 136 chemicals that reduce growth at 2 μM or less (Table S3). We then used the extended-connectivity fingerprint (ECFP) algorithm^72^ to convert all LATCA compound structures into numerical fingerprints. ECFPs were then used to compute structural similarity of pairs of compounds on a scale of 0 to 1 using Tanimoto coefficients^73^. The set of Tanimoto coefficients between all pairs of inhibitors was condensed into a usable network and visualized using Cytoscape^74^. We then used the strongest inhibitors to further reduce the similarity network to 28 clusters of structures exhibiting high levels of biological activity and selected 52 of these chemicals for subsequent treatment of the Chlamydomonas mutant library (Extended Data Fig. 1, Table S3, and Extended Data File 1).

### Library pooling and competitive growth experiments

The first two rounds of mutant library screening (R1, R2) were performed with the entire mutant collection (550 384-mutant array plates) in 20-liter carboys (Table S1 and Table S2). Mutants were pooled from five days old 384-colony array plates into liquid TAP medium at room temperature and low light. In R1, we pooled ten copies of eight plates (#668 to #670) in the collection to test how quantitatively we can track the relative abundance of mutants in the starting population. In R2, we pooled one set of the mutant collection (plates #597 to #670) from 384-colony array plates and another set from 1,536-colony array plates (#101 to #596) to test the performance of denser colony arrays for pooled screens.

Subsequent rounds of mutant library screening (R3-R6) were performed on the re-arrayed library (245 384-mutant array plates) in 2-liter bottles. Mutants were pooled from five days old 1,536-colony array plates. Condensing the library from 384 to 1,536-colony array plates helped to both homogenize colony growth and reduce the laborious pooling procedure.

We produced subpools each containing cells from eight 384 or 1,535-colony array plates by using sterile glass spreaders to pool cells the plates into 50 ml conical tubes containing 40 ml of TAP medium. These subpools were mixed by pipetting to break cell clumps using a 10 ml serological pipette with a P200 tip attached to it. Then, all subpools were combined into the final mutant collection pool by pipetting the subpools through a 100 μm cell strainer (VWR 10054-458). The final pool was mixed using a magnetic stir bar, and the cell density was measured (Countess, Invitrogen) and adjusted to 1×10^5^ cells ml^−1^. For experiments not performed in TAP medium, cells were washed twice with the actual medium used for pooled screens after pelleting (1000x g, 5 min, room temperature).

Aliquots of 2×10^8^ cells were pelleted (1000x g, 5 min, room temperature) by centrifugation and frozen to determine the relative abundance of each mutant in the starting population. These samples are denoted as “Initial”.

Cultures were inoculated with 2×10^4^ cells ml^−1^ in transparent 20-liter carboy tanks (R1 and R2) or standard 2-liter bottles (R3 - R6) using aliquots of the final mutant pool. Cultures were grown under a broad variety of conditions (Table S2). Unless otherwise indicated, cells were grown in Tris-Acetate-Phosphate (TAP) medium with modified trace elements at pH 7.5 under constant light (100 μmol photons m^−2^ s^−1^) at 22 °C, aerated with air and mixed using a conventional magnetic stirrer at 200 rpm. The cell density of competitive growth experiments was tracked and aliquots of 2×10^8^ cells were pelleted by centrifugation after seven doublings, when the culture reached approximately 2×10^6^ cells ml^−1^. Cell pellets were frozen for subsequent DNA extraction and barcode quantification.

### DNA extraction

Total genomic DNA was extracted from frozen cell pellets representing 2×10^8^ cells of each sample (initial, control, and treatment).

First, frozen pellets were thawed at room temperature and resuspended in 1.6 ml 0.5x SDS-EB (1% SDS, 200 mM NaCl, 20 mM EDTA, and 50 mM Tris-HCl, pH 8.0).

Second, 2 ml of phenol:chloroform:isoamyl alcohol (25:24:1) was added to each sample and mixed by vortexing. This solution was then transferred into 15 ml Qiagen MaXtract High Density tubes (Cat No./ID: 129065) and centrifuged at 3,500 rpm for 5 minutes. Subsequently, the aqueous phase was transferred to a new 15 ml conical tube, 6.4 μl RNase A was added and the solution was incubated at 37 °C for 30 minutes. The phenol/chloroform: isoamyl alcohol extraction was then repeated, and the aqueous phase was transferred into a new 15 ml Qiagen MaXtract High Density tube before adding 2 ml chloroform: isoamyl alcohol. This solution was mixed by vortexing and centrifuged at 3,500 rpm for 5 minutes then, 400 μl aliquots of the aqueous phase were transferred to 1.5 ml reaction tubes for DNA precipitation.

Third, 1 ml of ice-cold 100% ethanol was added to the solution to precipitate DNA. The tubes were gently mixed and incubated at –20 °C overnight. The DNA was pelleted at 13,200 rpm and 4 °C. The supernatant was discarded and the pellet washed in 1 ml 70% ethanol. The supernatant was discarded again and the pellet was air-dried before resuspension in 50 μl water. Subsequently, the elution fractions of each sample were pooled and the DNA concentration was measured using a Qubit fluorometer (Invitrogen).

### Internal barcode amplification and Illumina library preparation

Internal barcodes were amplified using Phusion Hot Start II (HSII) DNA Polymerase (Thermo Fisher, F549L). Sequence information for all primers used in this study is summarized in Table S10.

The 50 μl PCR mixture for 5’ barcode amplification contained: 125 ng genomic DNA, 10μl GC buffer, 5 μl DMSO, 1 μl dNTPs at 10 mM, 1 μl MgCl2 at 50 mM, 2.5 μl of each primer at 10 μM, and 1 μl Phusion HSII polymerase. Eight tubes of the PCR mixture were processed per sample and incubated at 98 °C for three minutes, followed by ten three-step cycles (98 °C for 10 s, 58 °C for 25 s and 72 °C for 15 s), and then eleven two-step cycles (98 °C for 10 s, 72 °C for 40 s).

The 50 μl PCR mixture for 3’ barcode amplification contained: 125 ng genomic DNA, 10μl GC buffer, 5 μl DMSO, 1 μl dNTPs at 10 mM, 2 μl MgCl2 at 50 mM, 2.5 μl of each primer at 10 μM, and 1 μl Phusion HSII polymerase. Eight tubes of the PCR mixture were processed per sample and incubated at 98 °C for three minutes, followed by ten three-step cycles (98 °C for 10 s, 63 °C for 25 s and 72 °C for 15 s), and then eleven two-step cycles (98 °C for 10 s, 72 °C for 40 s).

The PCR products of each sample were pooled for further processing. First, successful PCR was confirmed on a TBE 8% agarose gel in 1 x Tris Borate EDTA before concentrating the PCR products on a Qiagen MinElute column and measuring the DNA concentration on a Qubit fluorometer. Second, 200-250 ng of up to 16 3’ or 5’ PCR products were combined into an Illumina HiSeq2000 library. Third, the internal barcode bands of the Illumina HiSeq2000 libraries were gel-purified and subjected to quality control on an Agilent Bioanalyzer. In addition, DNA concentration was determined on a Qubit fluorometer. Fourth, HiSeq2000 libraries were sequenced at the Genome Sequencing Service Center at Stanford University (3155 Porter Dr., Palo Alto, CA 94304).

### Data analysis

Initial reads were trimmed using cutadapt version 1.7.1^75^ using the command “cutadapt -a <seq> -e 0.1 -m 21 -M 23 input_file.gz -o output_file.fastq”, where <seq> is GGCAAGCTAGAGA for 5’ data and TAGCGCGGGGCGT for 3’ data. Barcodes were counted by collapsing identical sequences using “fastx_collapser” (http://hannonlab.cshl.edu/fastx_toolkit) and denoted as “_read_count”. Barcode read counts for each dataset were normalized to a total of 100 million and denoted as “_normalized_reads” (Table S11). Replicate control treatments performed in the same screening round were averaged by taking the mean of the normalized read counts to generate the average normalized read count and by summing the read counts to generate the average read count. Control treatments that were averaged are denoted with “average” and can be found in Table S12. Mutants in the library contain on average 1.2 insertions^12^, each of which may contain a 5’ barcode, a 3’ barcode, both barcodes, or potentially more than two barcodes if multiple cassettes were inserted at the loci. To represent a given insertion within a mutant, we selected a single barcode to represent it. All barcodes associated with the same gene and deconvoluted to the same library well and plate position were assumed to be from the same insertion and were then compared to identify the barcode with the highest read counts in the initial samples (R2-R6) to serve as the representative barcode.

To identify mutants with growth defects or enhancements due to a specific treatment, we compared the abundance of each mutant after growth under the treatment condition to its abundance after growth under a control condition. We called this comparison a “screen”, and the ratio of these abundances the “mutant phenotype”. In order for a phenotype to be calculated, we required the control treatment to have a read count above 50.

To identify high-confidence gene-phenotype relationships we developed a statistical framework that leverages multiple independent mutant alleles. For each gene, we generated a contingency table of the phenotypes, P, by counting the number of alleles that met the following thresholds: [P < 0.0625, 0.0625 ≤ P < 0.125, 0.125 ≤ P < 0.25, 0.25 ≤ P < 0.5, 0.5 ≤ P < 2.0, 2.0 ≤ P < 4.0, 4.0 ≤ P < 8.0, 8.0 ≤ P < 16.0]. Only alleles that were confidence level 4 or less, had an insertion in CDS/intron/5’UTR feature, and had greater than 50 reads in the control condition were included in the analysis. A p-value was generated for each gene by using Fisher’s exact test to compare a gene’s phenotype contingency table to a phenotype contingency table for all insertions in the screen. A false discovery rate was performed on the p-values of genes with more than 2 alleles using the Benjamini-Hochberg method^76^. To determine a representative phenotype for a gene, the median phenotype for all alleles of that gene that were included in the Fisher’s exact test was used. For some analysis these gene phenotypes were normalized by setting the median value of all gene phenotypes in a screen to zero. Clustering was performed with Python packages SciPy and Seaborn. Data in Fig. 4a was clustered using the ‘correlation’ metric and ‘average’ method for the linkage algorithm. Spearman and Pearson correlations were calculated in Pandas. Transcriptome correlation data was collected, curated, and analyzed in the Merchant laboratory. Data was plotted and visualized with the Python packages Matplotlib and Seaborn.

To determine if biological functions were associated with specific screens we performed a Gene Ontology (GO) term enrichment analysis. Using the same approach as with genes, we generated contingency tables of mutant phenotypes for each GO term. If a mutant’s insertion is within a gene that had multiple GO term annotations, the mutant’s phenotype data was added to each GO term’s contingency table. A p-value was generated for each GO term by using Fisher’s exact test to compare a GO term’s phenotype contingency table to a phenotype contingency table for all GO terms in the screen. A false discovery rate was performed on the p-values using the Benjamini-Hochberg method^76^. Clustering were performed in the

All analysis was performed using JGI Phytozome release v5.0 of the Chlamydomonas assembly and v5.6 of the Chlamydomonas annotation^77^.

### Immunoblot materials IDA5, NAP1

Cells were collected by centrifugation, frozen in liquid nitrogen, and subsequently resuspended in 100 μl of ice-cold PNE buffer (10 mM phosphate pH 7.0, 150 mM NaCl2, 2 mM EDTA) supplemented with a complete protease-inhibitor cocktail (Roche; 11697498001) and disrupted by vortexing with acid-washed glass beads. In some experiments using anti-actin antibodies, these samples were mixed directly with SDS-PAGE sample buffer, boiled for 3 min, and cleared of debris by centrifugation at 12,000 x g for 10 min at 4 °C before electrophoresis. In all other experiments, 100 μl of PNE buffer + 2% NP-40 were added to the samples after cell disruption, and the samples were then incubated for 10 min on ice and cleared by centrifugation at 12,000 x g for 10 min at 4 °C before adding SDS-PAGE sample buffer. We did not observe any difference in abundance or solubility of IDA5 or NAP1 between the two methods. SDS-PAGE was performed using 11% Tris-glycine (for IDA5 and NAP1). Blots were stained using a mouse monoclonal anti-actin antibody (clone C4, EMD Millipore, MAB1501), which recognizes IDA5 but not NAP1; a rabbit antiNAP1 antibody (generous gift from Ritsu Kamiya and Takako Kato-Minoura), which recognizes NAP1 but not IDA5. HRP-conjugated antimouse-IgG (ICN Pharmaceuticals; 55564) or anti-rabbit-IgG (Southern Biotech; 4050-05) were used as secondary antibodies. Figures showing blots are cropped to show only the molecular weight ranges of interest: for IDA5 and NAP1, 37-50 kDa.

### MMS growth assays and VIPP2 immunoblot analysis

The following strains were used^17^: WT = CC-4533; *mars1* = *mars1-3*; *mars1:MARS1-D* = *mars1-3* transformed with the *MARS1-D* transgene containing a 3x-Flag epitope after Met139; *mars1:MARS1-D KD* = *mars1-3* transformed with a catalytically-inactive *MARS1-D* bearing the kinase active site D1871A mutation. Prior to starting liquid cultures in TAP media, all strains were restreaked in fresh TAP plates and grown in similar light conditions (i.e., ~50–70 μmol photons m^−2^ s^−1^, ~22 °C) for about 5-6 days. Prior to starting the MMS treatment, all strains were pre-conditioned in liquid cultures for about 3-4 days. Next, cell cultures were equally diluted to ~5 μg chlorophyll ml^−1^ and incubated in the presence or absence of MMS for 48 hours. A 1% (vol/vol) MMS stock solution (Sigma Aldrich # 129925) was freshly prepared in ddH20 at the beginning of each experiment. This MMS stock solution was further diluted 200 times directly into TAP media to a final concentration of 0.05% (vol/vol). All chlorophyll concentration measurements were performed using a previously described methanol extraction method^78^.

VIPP2 and alpha-TUBULIN immunoblot analyses were carried out as described^17^, using denatured total protein samples prepared from liquid cultures incubated for 27 hours in the presence or absence of 0.05% (vol/vol) MMS.

### Cilia and LatB-related Chlamydomonas experiments

Mutants used in this study are listed in Table S13. Individual mutants were grown with gentle agitation at 100 μmol photons m^−2^ s^−1^. Disruption of *LAT5*, *LAT6*, and *LAT7* genes (Cre17.g721950, Cre15.g640101, and Cre11.g482750) in the original isolates of *lat5-1*, *lat6-1*, *lat7-2* were confirmed by PCR. These mutants were then backcrossed with CC-124 or CC-125 three times, with perfect linkage of paromomycin resistance and LatB sensitivity in at least 10 tetrads confirmed after each round. The backcrossed strains and the previously established *lat1-5*, *lat2-1*, *lat3-1*, and *nap1-1* mutants in the CC-124 background^68^ were spotted on TAP agar containing 0.1% DMSO with or without 3 μM LatB as 5x serial dilutions.

Cilia mutants were grown in liquid TAP medium until they reached exponential phase. Cells were then mounted in u-Slide 8-well chambers (Ibidi, 80826) with 2% low melting point agarose (Sigma, A9414). Cilia defects were scored using a Leica DMi8 inverted microscope. Cilia length was measured using Fiji. Cilia swimming behavior was scored using TAP agar plates with 0.15% agar. Latrunculin B (Sigma, L5288) treatments were performed on TAP agar plates supplemented with 3 μM LatB and spotted in 10-fold serial dilutions.

### Arabidopsis experiments

Mutants used in this study are listed in Table S13. Seeds were surface-sterilized in 20% bleach for 5 min. Seeds were then rinsed with sterile water four times and stored at 4 °C for three days in the dark. After stratification, seeds were sown into square 10 cm x 10 cm petri plates containing full strength Murashige and Skoog (MS) medium (MSP01-50LT), 1% agar (Duchefa, 9002-18-0), 1% sucrose, 0.05% MES, and adjusted to pH 5.7 with 1 M KOH. Seedlings were grown in the presence of LatB (Sigma, L5288) or mock control containing equivalent volume of the LatB solvent, DMSO. Plates were imaged using a CanonScan 9000 flatbed scanner. Root lengths were quantified using Fiji. Two-way ANOVA and data visualization were done using Python.

